# Slow Ionic Dynamics Reshape Bistability in a Biophysical Mean-Field Model of Brain Activity

**DOI:** 10.64898/2026.10.22.760893

**Authors:** Sourin Chatterjee, Charlotte Maschke, Spase Petkoski, Huifang Wang, Damien Depannemaecker, Viktor Jirsa

## Abstract

Brain states such as sleep, seizures, and resting wakefulness are often described in terms of transitions between distinct population states. Biophysically inspired mean-field models that couple neuronal firing statistics to slow ionic homeostasis provide a mechanistic bridge between detailed single-neuron physiology and macroscopic brain activity. Here we examine this issue in a biophysical meanfield model of cortical populations driven by potassium exchange and compare its dynamical repertoire with a matched spiking network across six physiological and pathological regimes. The fast subsystem contains a substantial region in which low- and high-activity fixed points coexist, but this conditional bistability is not inherited by the physiological resting branch of the complete slow-fast system. A generic fast-slow calculation identifies how feedback through a dynamical ionic variable changes the stability conditions of the fast subsystem, and an exact equilibrium reduction of the KIonEx-MF equations yields a single equilibrium on the physiological resting branch over the range examined. This resting state is excitable: perturbations generate intermittent synchronized events in both the mean-field and spiking-network descriptions. The results show how slow ionic dynamics can reshape conclusions drawn from a frozen fast subsystem and provide noise-driven excitability as an alternative route to spontaneous population transitions.

## 1 Introduction

Understanding how population-level brain dynamics emerge from interactions among neurons is a central problem in computational neuroscience. Detailed spiking-neuron models resolve cellular excitability and synaptic transmission, but their dimensionality and computational cost make them difficult to deploy at whole-brain scale or to relate directly to macroscopic measurements such as electroencephalography (EEG) and functional magnetic resonance imaging (fMRI). Mean-field models provide a practical bridge between these scales by replacing large populations of interacting neurons with a small number of collective variables while retaining links to the underlying physiology [1, 2]. Such models have been used to study resting-state dynamics [3, 4], large-scale oscillations [5], seizure dynamics [6, 7], neuropsychiatric disorders [8], and spontaneous state transitions [9] (see [10] for a review).

Modeling seizure dynamics is of substantial clinical interest because a mechanistic account of seizure generation, propagation and termination can inform the diagnosis and treatment of drug-resistant epilepsy [11]. Mean-field models of seizures range from phenomenological descriptions, in which the equations are chosen to reproduce observed dynamics, to biophysically inspired formulations whose variables map onto identifiable cellular processes. Phenomenological models such as the Epileptor [6, 7] reproduce a broad range of epileptic dynamics and have been used to identify epileptogenic regions, but their state variables do not have a one-to-one correspondence with underlying biophysical processes [12, 13]. Ion-based models provide a complementary route by representing quantities that can be connected more directly to cellular physiology and experimental manipulation [14]. Slow ionic processes are particularly important in pathological transitions such as seizures and spreading depolarization [15–19]. Extracellular potassium evolves much more slowly than membrane voltage through diffusion, Na^+^/K^+^ pumping and glial buffering [20, 21], and its accumulation is a well-established contributor to seizure-like transitions and spreading depolarization [22–26].

Alongside epilepsy, resting-state dynamics has become a major target of mean-field modeling because spontaneous brain activity remains spatially and temporally structured even in the absence of an explicit task [27]. Classical neural-mass frameworks, including the Wilson-Cowan [28] and Jansen-Rit [29, 30] models, and more recent Hopf normal-form whole-brain models [31, 32], provide low-dimensional descriptions of this population activity. Montbrió et al. [33] derived exact macroscopic equations for an infinite heterogeneous population of quadratic integrate-and-fire neurons. Depending on its parameters, this reduced system can support coexisting lowand high-activity states. Bistable local dynamics therefore provides one route by which fluctuations can generate transitions between population states, related to the up-down dynamics observed in cortical recordings [34–36]. Related mean-field models embedded in whole-brain networks can reproduce important features of resting-state dynamics [37, 38].

Depannemaecker et al. [12] showed that a Hodgkin-Huxley-inspired neuron model coupled to ion dynamics reproduces spiking, tonic firing, bursting, seizure-like events, sustained activity and depolarization block within a single biophysical framework. This builds on earlier work showing that elevated extracellular potassium can render Hodgkin-Huxley-type neurons bistable [39–41]. The single-neuron formulation subsequently informed a population-level mean-field model driven by ion exchange [42]. The resulting model contains a fast subsystem describing population firing, membrane potential and gating dynamics, together with slower variables describing ionic exchange. Here we examine how these two timescales shape the model’s dynamical repertoire and, in particular, whether attractor structure identified with the ionic variables fixed persists once those variables are allowed to evolve.

We first characterize the potassium-ion-exchange mean-field (KIonEx-MF) model across its two principal control parameters and compare the resulting regimes with simulations of the matched spiking network. We then use a slow-fast decomposition to examine the fixed-point structure of the model. The fast subsystem contains a substantial region in which two stable fixed points coexist, but this conditional bistability does not by itself establish bistability of the complete system. We therefore examine explicitly how the slow potassium dynamics changes the stability structure, first in a generic fast-slow model and then in the KIonEx-MF equations. Finally, we return to the low-activity physiological regime and show that the deterministic fixed point is excitable. This provides a candidate mechanism for spontaneous population events in which fluctuations generate transient excursions without requiring switching between two deterministic stable states.

## 2 Results

### 2.1 Dynamical Repertoire of the Model and SNN Comparison

To understand how extracellular potassium homeostasis and recurrent synaptic coupling shape the collective dynamics of the model, we systematically explored the effects of two parameters: extracellular potassium concentration (*K*_*bath*_) and recurrent synaptic strength (*J*). These two parameters regulate complementary aspects of cortical excitability. *K*_*bath*_ is directly related to astrocytic buffering of extracellular potassium, and impairments in this process have been associated with several pathological conditions. The parameter *J*, on the other hand, determines the strength of recurrent synaptic interactions and can be altered by synaptic loss.

The parameter sweep revealed six qualitatively distinct dynamical regimes, associated with six electrophysiological patterns. To identify these regimes, we quantified six summary statistics of the membrane-potential time series: the standard deviation, median, minimum and maximum of *V*, the number of detected population-voltage peaks, and the mean peak height. Thresholds applied jointly to these statistics defined the regime boundaries (red lines in Fig. 1(a)); the corresponding parameter maps are shown in the *SI Appendix*, Fig. S1. We repeated the classification from multiple initial conditions to check that the identified regimes were not specific to a single starting state.

**Fig. 1.**
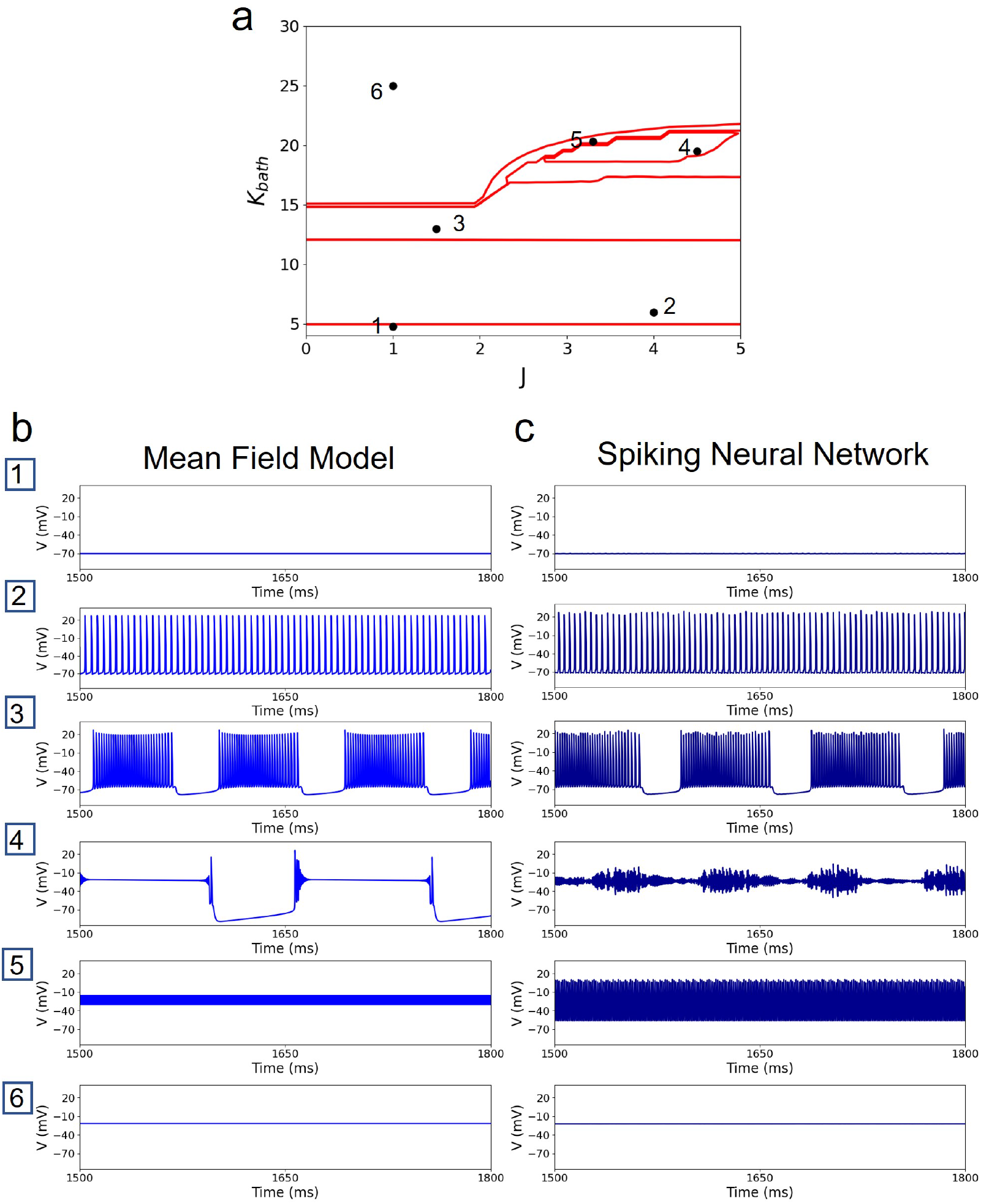
Dynamical repertoire of the model. (a) Parameter sweep across the extracellular potassium bath concentration (*K*_*bath*_) and recurrent synaptic strength (*J*) plane, delineating six qualitatively distinct dynamical regimes (boundaries shown in red, derived from the membrane-potential summary statistics shown in *SI Appendix*, Fig. S1). Representative parameter values for each regime are marked and numbered 1-6. (b) Membrane potential (*V*) time series of the mean-field (MF) model for the six representative regimes: (1) asynchronous activity, (2) synchronized population firing, (3) epileptic-like activity, (4) depolarization-recovery activity, (5) rapid synchronized firing, and (6) depolarization block. (c) Corresponding population-averaged membrane potential from the spiking neural network (SNN), simulated with matched parameters, shown for qualitative comparison with (b). Raster plots of SNN can be found in *SI Appendix*, Fig. S2.

We labeled the six regimes as: (1) asynchronous activity, (2) synchronized population firing, (3) epileptic-like activity, (4) depolarization-recovery activity, (5) rapid synchronized firing, and (6) depolarization block, as illustrated in Fig. 1(b). Together, these regimes span a broad range of network behaviors, from stable resting activity to persistent pathological depolarization. Representative values of *K*_*bath*_ and *J* were selected to illustrate each regime. During asynchronous activity, the mean-field membrane potential remains near −70.3 mV, corresponding to a stable resting state. Increasing excitability leads to synchronized population firing, with recurrent excursions from approximately −71.9 mV to +27.0 mV. In the epileptic-like regime, the model exhibits a period of rapid oscillatory activity before spontaneously terminating. Depolarization-recovery activity alternates between a depolarized epoch and recovery toward more hyperpolarized voltages. Rapid synchronized firing consists of high-frequency population oscillations, while in the depolarization-block regime the mean-field membrane potential remains near −21.8 mV without recovery.

To determine whether these dynamical regimes are preserved beyond the mean-field description, we performed simulations of the corresponding spiking neural network (SNN). The same combinations of *K*_*bath*_ and *J* were used for all representative cases except Case 4, for which the SNN required a shifted value *K*_*bath*_ = 22.6 to obtain depolarization-recovery dynamics. Previous studies have shown that the correspondence between spiking and mean-field models is not always exact: a dynamical regime observed in the mean-field model at a particular parameter value may appear in the spiking network at a slightly shifted parameter value [42]. For asynchronous activity, synchronized firing, epileptic-like activity and depolarization block, the SNN reproduces the qualitative dynamics observed in the mean-field model. Larger differences occur in Cases 4 and 5. In Case 4, the SNN shows both a smaller voltage excursion and less regular phase durations than the mean field; in Case 5, the oscillation amplitude and population-peak count differ substantially between the two descriptions. These two cases expose limitations of the mean-field approximation, whereas the remaining four regimes show close qualitative and, where quantified, quantitative correspondence.

### 2.2 Understanding Stability of the Fixed Points

To identify the dynamical mechanisms underlying the observed activity regimes, we analyzed the fast subsystem of the model using a slow-fast decomposition. Since the slow variables (Δ*K*_*i*_ and *K*_*g*_) evolve on a much longer timescale than the membrane potential (*V*), recovery variable (*x*) and gating variable (*n*), they can be treated as quasi-static parameters during the analysis. For each representative regime, we computed the three nullclines of the fast subsystem and overlaid the simulated trajectories in the three-dimensional phase space (Fig. 2). For stationary cases, the slow variables were fixed at their asymptotic values. In Cases 3, 4, and 5, however, the slow variables themselves oscillate (*SI Appendix*, Figs. S3 and S4); representative values along the corresponding slow-variable cycle were therefore used for the fast-subsystem nullcline analysis.

**Fig. 2.**
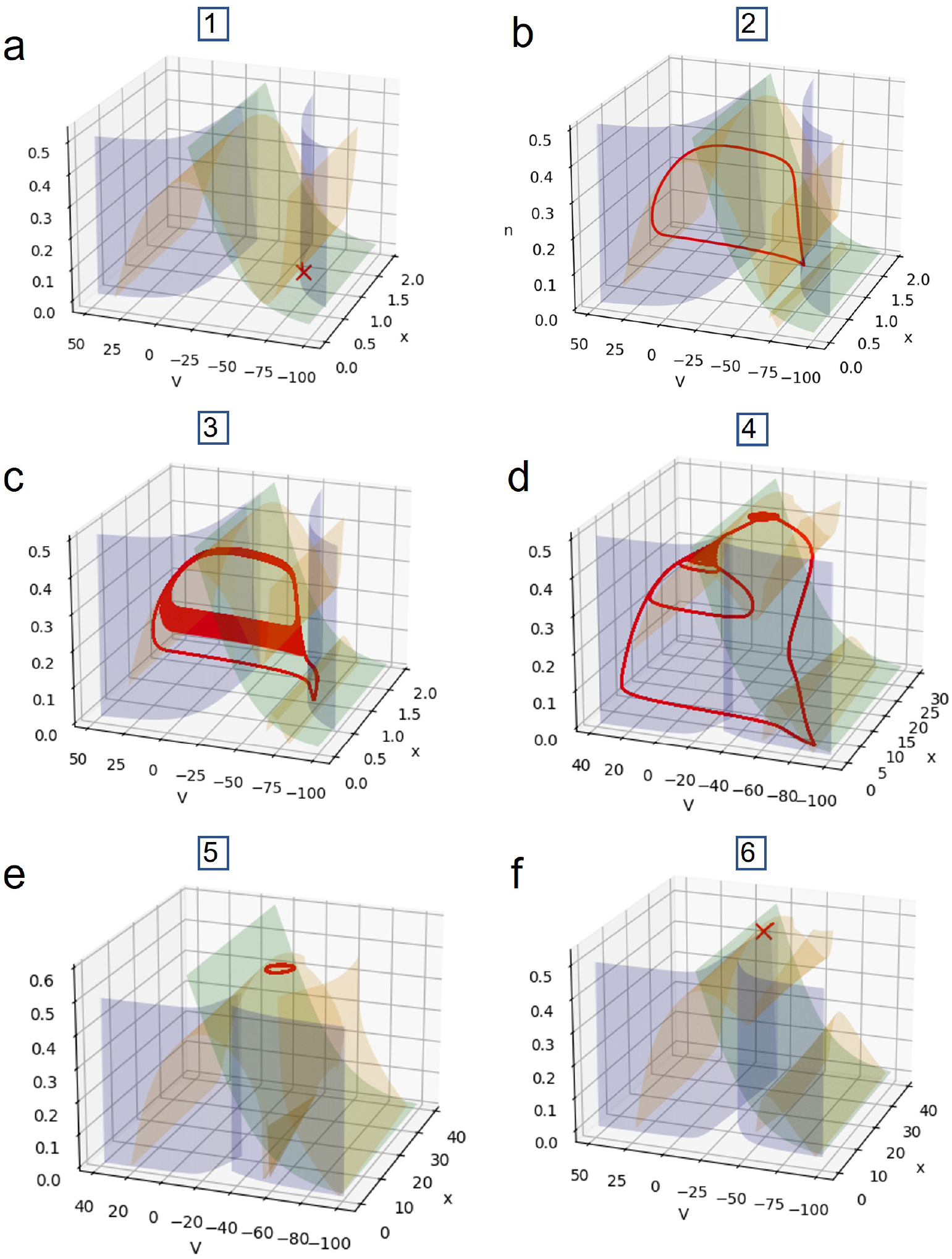
Fast-subsystem phase portraits underlying the six dynamical regimes. For each representative case in Fig. 1, the slow variables (Δ*K*_*i*_, *K*_*g*_) were fixed at their simulated steady-state (or a representative intermediate) value and treated as quasi-static parameters of the fast subsystem (*x, V, n*). Colored surfaces show the nullclines of *x* (blue), *V* (orange) and *n* (green); and the red curve shows the simulated fast-variable trajectory from Fig. 1, overlaid on the nullcline geometry. (a) Case 1, asynchronous activity: a stable fixed point coexists with an unstable fixed point. (b) Case 2, synchronized population firing: a stable limit cycle. (c) Case 3, epileptic-like activity. (d) Case 4,depolarization-recovery activity. (e) Case 5, rapid6synchronized firing: a stable limit cycle. (f) Case 6, depolarization block: a stable depolarized fixed point together with an unstable fixed point. For Cases 3, 4, and 5, variations in the slow parameters are visualized in the *SI Appendix*.

The phase-space analysis provides a mechanistic explanation for the different dynamical regimes. During asynchronous activity (Case 1), the system exhibits a stable fixed point together with an additional unstable fixed point. Meanwhile, synchronized population firing is characterized by a stable limit cycle. The epileptic-like and depolarization-recovery regimes display more complex trajectories, reflecting transitions between two distinct activity states, and therefore cannot be described by a single stable fixed point. Rapid synchronized firing also exhibits a limit-cycle solution, whereas depolarization block corresponds to a stable depolarized fixed point accompanied by an unstable fixed point.

Although both asynchronous activity and depolarization block correspond to stationary states, their responses to perturbations are markedly different. To investigate this difference, we applied a small depolarizing pulse (2 mV) to the membrane potential and examined the resulting transient dynamics. In the asynchronous regime, this perturbation generates a large transient excursion before the system relaxes back to the resting state (Fig. 3, left), a characteristic signature of an excitable fixed point. In contrast, the same perturbation produces only a slight response and returns to its state very quickly during depolarization block (Fig. 3, right), consistent with the loss of neuronal responsiveness in this regime. We further examined the separatrix associated with these fixed points and found that relatively small perturbations are sufficient to trigger large excursions in the asynchronous state, whereas substantially larger perturbations are required during depolarization block. These responses identify the asynchronous regime as an excitable fixed point.

**Fig. 3.**
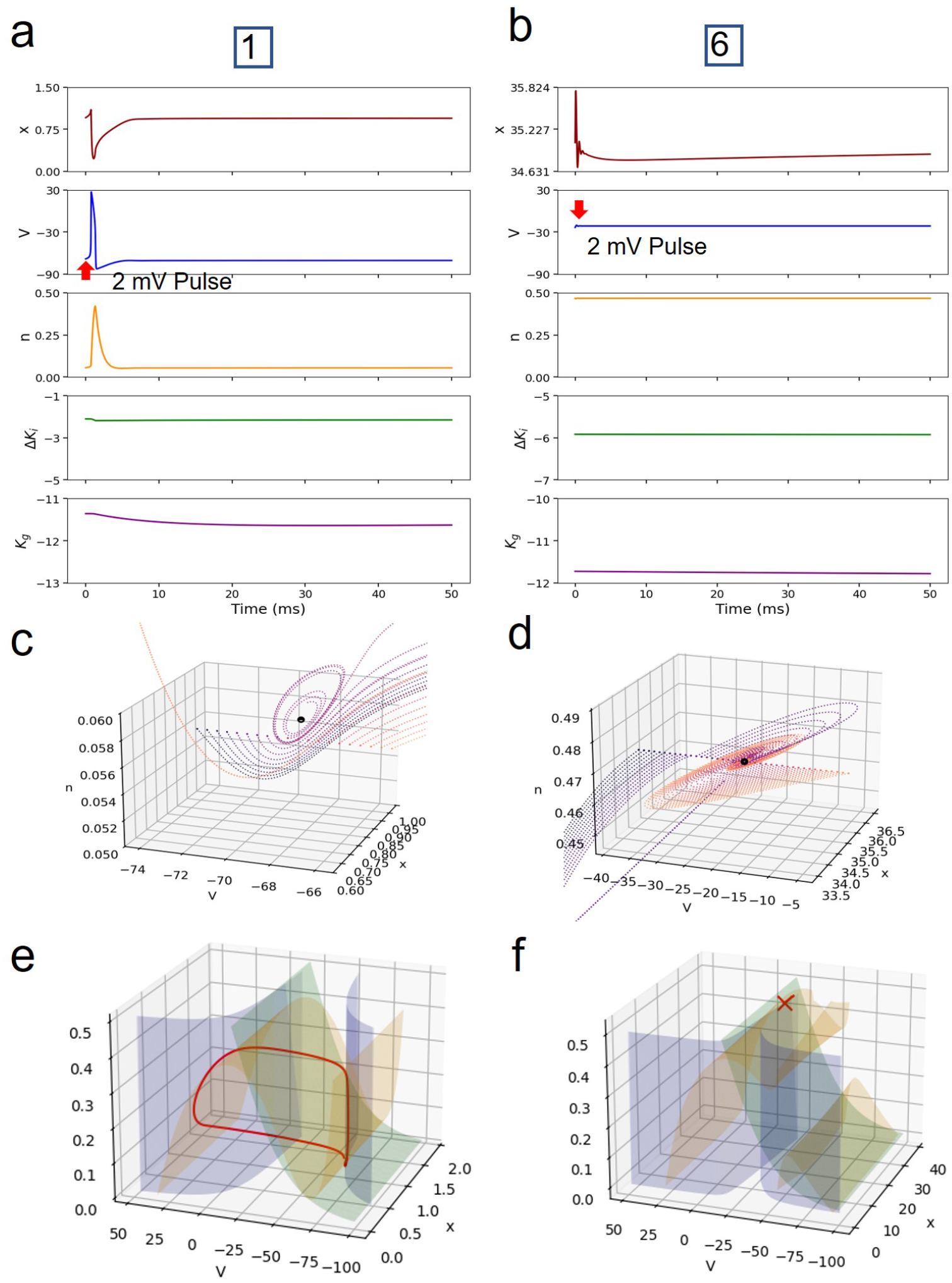
Excitability revealed by a depolarizing perturbation. A 2 mV depolarizing pulse (red arrow) was applied to the membrane potential in the asynchronous regime (Case 1, a, c, e) and the depolarization block regime (Case 6, b, d, f). (a, b) Time series of *x, V, n*, Δ*K*_*i*_ and *K*_*g*_ following the perturbation, for Case 1 (a) and Case 6 (b). (c, d) Fast-subsystem trajectory in (*V, x, n*) space immediately after the perturbation, for Case 1 (c) and Case 6 (d). (e, f) The same trajectory (red) overlaid on the fast-subsystem nullcline geometry (as in Fig. 2), together with the pre-perturbation fixed point(s), for Case 1 (e) andCase 6 (f). The asynchronous state (a, c, e) shows a large transient excursion before relaxing back to rest, characteristic of an excitable fixed point, whereas the depolarization block state (b, d, f) shows only a small, rapidly damped transient before returning to its depolarized equilibrium.

### 2.3 The Fast Subsystem Dynamics

The nullcline analysis in Fig. 2 characterizes the fast subsystem only at the six pairs of slow-parameter values (Δ*K*_*i*_, *K*_*g*_) obtained by our representative cases. To get a more complete picture, we performed a parameter sweep of Δ*K*_*i*_ and *K*_*g*_ over a two-dimensional grid and classified the fast subsystem’s asymptotic behavior at each point, yielding the nine qualitative classes shown in Fig. 4, together with a representative 20 ms example trace of *V* for each (bottom, panels 1-8). In order of increasing dynamical complexity, these are: a single low-activity fixed point (class 1); a single high-activity fixed point (class 2); coexistence of both (class 3); a single largeamplitude limit cycle (class 4); coexisting largeand small-amplitude limit cycles (class 5); a low-activity fixed point coexisting with a large limit cycle (class 6); a high-activity fixed point coexisting with a large limit cycle (class 7); a highand a low-activity fixed point coexisting with a large limit cycle (class 8); and parameter combinations for which the fast-subsystem equations are numerically ill-defined (class 9, Failure), marked separately.

**Fig. 4.**
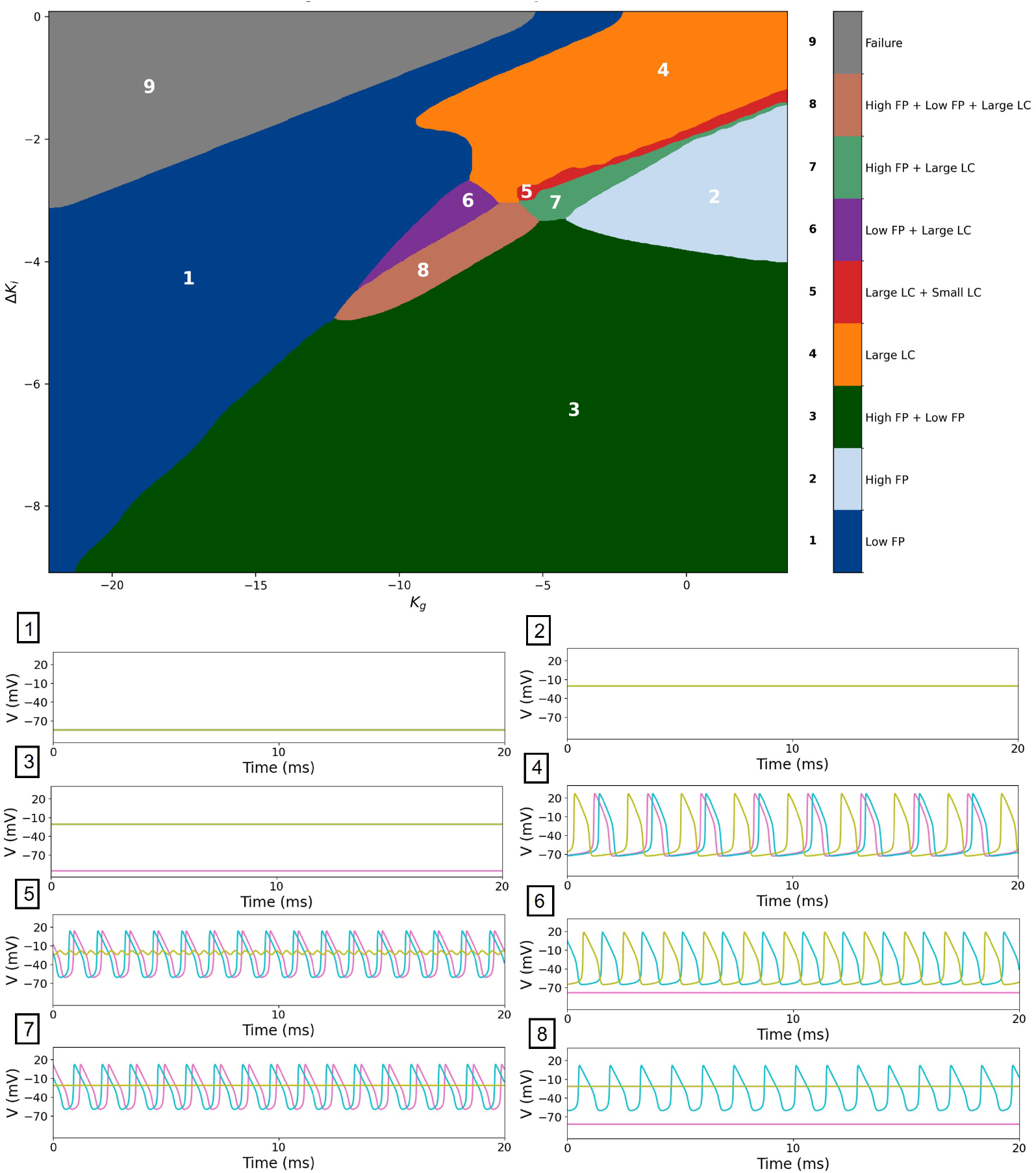
Classification of fast-subsystem dynamics across the (Δ*K*_*i*_, *K*_*g*_) parameter plane. For each grid point, Δ*K*_*i*_ and *K*_*g*_ were held fixed and the fast subsystem (*x, V, n*) was simulated to identify its asymptotic behavior, yielding nine qualitatively distinct classes (numbered and colorcoded 1-9; legend at right). Below, representative 20 ms time series of *V* are shown for each class after transient (panels 1-8, colored as in the map), each simulated from three distinct initial conditions (*x*_0_, *V*_0_, *n*_0_), chosen to sample different basins of attraction of the fast subsystem: (1.08, *−* 68.20, 0.04) (magenta), (1.32, *−* 62.17, 0.18) (cyan), and (24.28, *−*21.74, 0.46) (olive). Class 9 (Failure) denotes parameter combinations for which the fast-subsystem equations become numerically ill-defined, as the argument of the logarithmic terms in the ionic currents takes a negative value for that combination of Δ*K*_*i*_ and *K*_*g*_. **Abbreviations:** FP, fixed point (Low, low-activity; High, high-activity); LP, limit cycle (Small, small-amplitude; Large, large-amplitude).

A substantial region of this map, class 3 shown in green (Fig. 4), contains two stable fixed points. For points in this region, independent initial conditions of the fast subsystem converge to lowand high-activity fixed points, respectively (see Methods). So from this analysis, a trajectory ending in class 3 would be expected to retain access to both stable branches. We tested this interpretation by overlaying the slow-variable trajectory (Δ*K*_*i*_(*t*), *K*_*g*_(*t*)) of each of the six representative cases of Fig. 1 on the classification map (*SI Appendix*, Fig. S5), marking the start and end of each trajectory in slow-variable space.

Two cases make clear why the frozen fast-subsystem map cannot be read as a bifurcation diagram of the complete system. In Case 5, the slow variables trace a closed orbit while the instantaneous frozen subsystem lies predominantly in a region classified as a low-activity fixed point; the full system therefore exhibits rapid oscillations rather than settling to that frozen attractor. In Case 6, the terminal (Δ*K*_*i*_, *K*_*g*_) values lie in the frozen bistable region, but the low-activity fast-subsystem fixed point at those slow-variable values does not satisfy the slow equilibrium condition 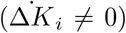 and is therefore not an equilibrium of the full five-dimensional system. Across the initial conditions tested, the complete system converges to the high-activity branch. The frozen map thus identifies conditional fast-subsystem attractors, whereas a full-system equilibrium must simultaneously satisfy both the fast and slow nullclines. We next ask whether two such equilibria exist on the physiological resting branch.

### 2.4 Effect of the slow ionic dynamics on bistability

#### 2.4.1 Generic fast-slow model

To determine whether the loss of bistability depends on the particular nonlinearities of the KIonEx-MF model, we first consider a minimal fast-slow system with the same local coupling structure. Near the cusp of the quadratic integrate-and-fire mean-field equations, the dynamics along the center direction can be reduced to the cubic normal form

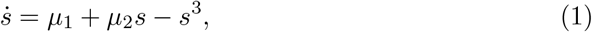

where *µ*_1_ and *µ*_2_ are unfolding parameters (see *SI Appendix*). For fixed *µ*_1_ and *µ*_2_, Eq. (1) contains the usual region in which two stable outer fixed points coexist with an unstable middle fixed point.

For the stability calculation we shift and rescale the cusp coordinates so that the uncoupled fast term is *f* (*s*) = *s* −*s*^3^ and write the intrinsic slow term as *q*(*u*) = *u* −*u*_0_.

The linearized potassium-current contribution is parameterized as

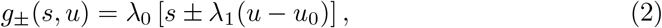

with *λ*_0_ *>* 0 setting the scaled fast-current coupling and *λ*_1_ *>* 0 the magnitude of the slow-variable contribution. The minus sign is used for the extracellular case (*g*_*u*_ *<* 0) and the plus sign for the intracellular case (*g*_*u*_ *>* 0). The coupled normal form is therefore

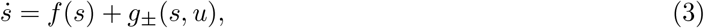

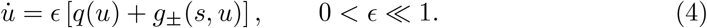

The complete algebra, including the fixed-point relations, Jacobian, and stability inequalities, is given in the *SI Appendix*. In the slow-fast limit, allowing *u* to evolve adds a stability condition that is absent when it is treated as a fixed parameter. We consider the two coupling signs separately.

##### Extracellular potassium

If the slow variable represents extracellular potassium while intracellular potassium is taken as approximately fixed, the linearized Nernst contribution gives

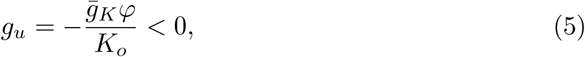

with *φ* = *RT/F*. In the slow-fast limit, stability of both outer branches requires the dimensionless coupling parameter *λ*_1_ to exceed

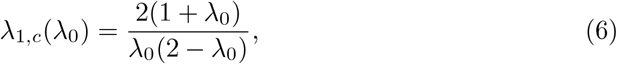

for 0 *< λ*_0_ *<* 2. The minimum value of this threshold is

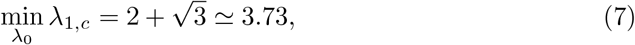

attained at

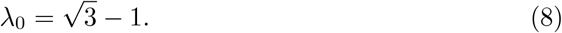

We can compare this condition with the coupling generated by the Nernst term over the range of potassium concentrations considered here. Defining

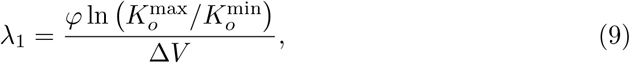

and taking *K*_*o*_≃4.8–16 mM together with a voltage excursion of approximately 65 mV gives

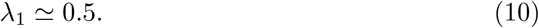

This remains well below the minimum value required by Eq. (6). Within this approximation, extracellular-potassium feedback is therefore too weak to preserve the two stable branches over the concentration range considered here.

##### Intracellular potassium

If the slow variable instead represents intracellular potassium, the sign of the coupling changes:

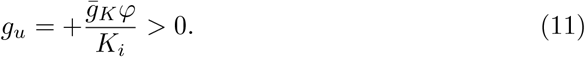

Repeating the stability calculation gives the conditions

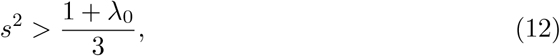

and

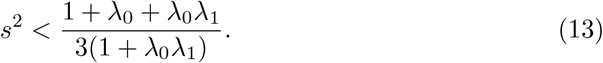

For these two intervals to overlap one would require

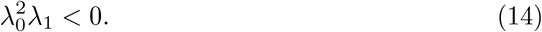

Since *λ*_0_ *>* 0 and *λ*_1_ *>* 0 by construction, this condition cannot be satisfied. Thus, within this reduced system, intracellular-potassium feedback does not permit the two fast-attracting branches to remain stable simultaneously. The two cases therefore impose different restrictions: extracellular feedback falls below the required stability threshold over the range examined here, whereas in the intracellular case the two stability intervals do not overlap.

When the stable branches do not exist, the coupled fast-slow system can instead develop small-amplitude oscillations close to a fold. The calculation in the *SI Appendix* yields the characteristic frequency scale

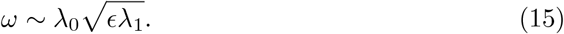

This square-root scaling arises from the interaction of the small fast eigenvalue near the fold with the explicit slow timescale.

#### 2.4.2 Bistability in the KIonEx-MF model

We next return to the KIonEx-MF model and ask whether two stable equilibria survive over the physiological resting range. At equilibrium of the *K*_*g*_ equation,

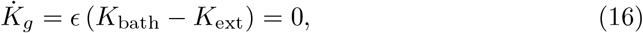

and therefore

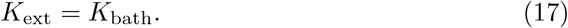

At an equilibrium, the *K*_*g*_ condition fixes extracellular potassium at the bath value and the remaining ionic degree of freedom is Δ*K*_*i*_. The other stationary conditions give

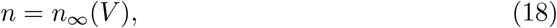

together with the branchwise algebraic relation obtained from 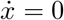for the firing rate. These relations are exact at a fixed point; no adiabatic approximation is required to determine equilibrium existence or location. They reduce the equilibrium problem to two algebraic equations in (*V*, Δ*K*_*i*_), with *K*_bath_ as an external parameter. Stability of any resulting equilibrium must still be determined from the full five-dimensional Jacobian (see *SI Appendix*).

The derivative of the Δ*K*_*i*_ equation with respect to Δ*K*_*i*_ contains the contributions

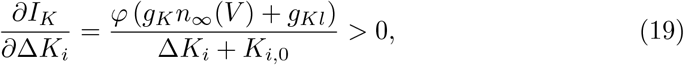

and

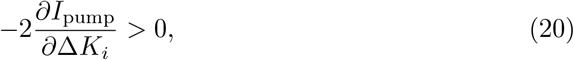

over the admissible range considered here. The corresponding Δ*K*_*i*_-nullcline of the reduced equations is therefore monotonic on this branch.

We solved this nullcline numerically for Δ*K*_*i*_ as a function of *V*, substituted the result into the *V* -nullcline and counted the resulting zero crossings while varying *K*_bath_. Only one crossing was found throughout the physiologically relevant resting branch examined here. Because the equilibrium elimination is exact, this establishes that two full-system equilibria cannot coexist on that resting branch over the range tested; it does not by itself constitute a stability calculation or an exhaustive count on the depolarized branch.

An independent check based directly on the sign of the product of the fast and slow coupling derivatives gives the same result over this range. Once *K*_*g*_ has relaxed and *K*_ext_ is constrained by *K*_bath_, the remaining feedback between *V* and Δ*K*_*i*_ has the same sign as the intracellular-potassium case of the generic calculation above.

We also searched for bistability while varying several additional model parameters, including *η*, Δ, *τ*_*n*_, *ϵ, γ* and *C*_*m*_, but did not identify a second stable fixed point over the parameter ranges examined.

The reduction above is exact for equilibrium existence because it is obtained by imposing 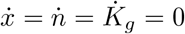 algebraically. Its limitations are instead twofold. First, the two-dimensional elimination does not preserve the eigenvalue spectrum, so stability and local bifurcations must be assessed from the full five-dimensional Jacobian. Second, the numerical root count reported here was carried out on the physiological resting branch. The branchwise firing-rate relation remains real on the depolarized branch as well (*SI Appendix*), but we do not use the resting-branch count as an exhaustive continuation of all depolarized equilibria.

### 2.5 Excitability as a Working Point for Network-Level Simulation

The absence of bistability in the physiological range leaves open the question of how spontaneous transitions could arise when this model is used as a node in a larger brain network. The asynchronous state provides one possibility. As shown in Fig. 3, it is a stable fixed point, but a sufficiently large perturbation produces a pronounced excursion before the trajectory returns to rest. We therefore tested whether stochastic fluctuations can generate such excursions without changing the deterministic fixedpoint structure.

We introduced stochastic perturbations into the membrane-potential and gating-variable equations. For additive noise,

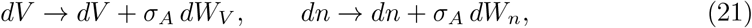

whereas for multiplicative noise,

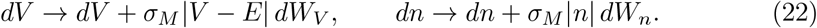

Here *dW*_*V*_ and *dW*_*n*_ denote independent Wiener increments.

For both noise implementations, weak fluctuations produced occasional large population events separated by much longer intervals during which the system remained close to the asynchronous fixed point (Fig. 5a). The corresponding spiking network showed the same qualitative behavior (Fig. 5b,c): firing remained irregular between events, whereas individual excursions produced transient recruitment of a substantial fraction of the population. We next varied the noise amplitude and counted the resulting population events. Events were rare or absent at low noise amplitudes and became progressively more frequent once fluctuations were large enough to drive the trajectory away from the resting state (*SI Appendix*, Fig. S6). The same qualitative dependence was obtained for additive and multiplicative noise.

**Fig. 5.**
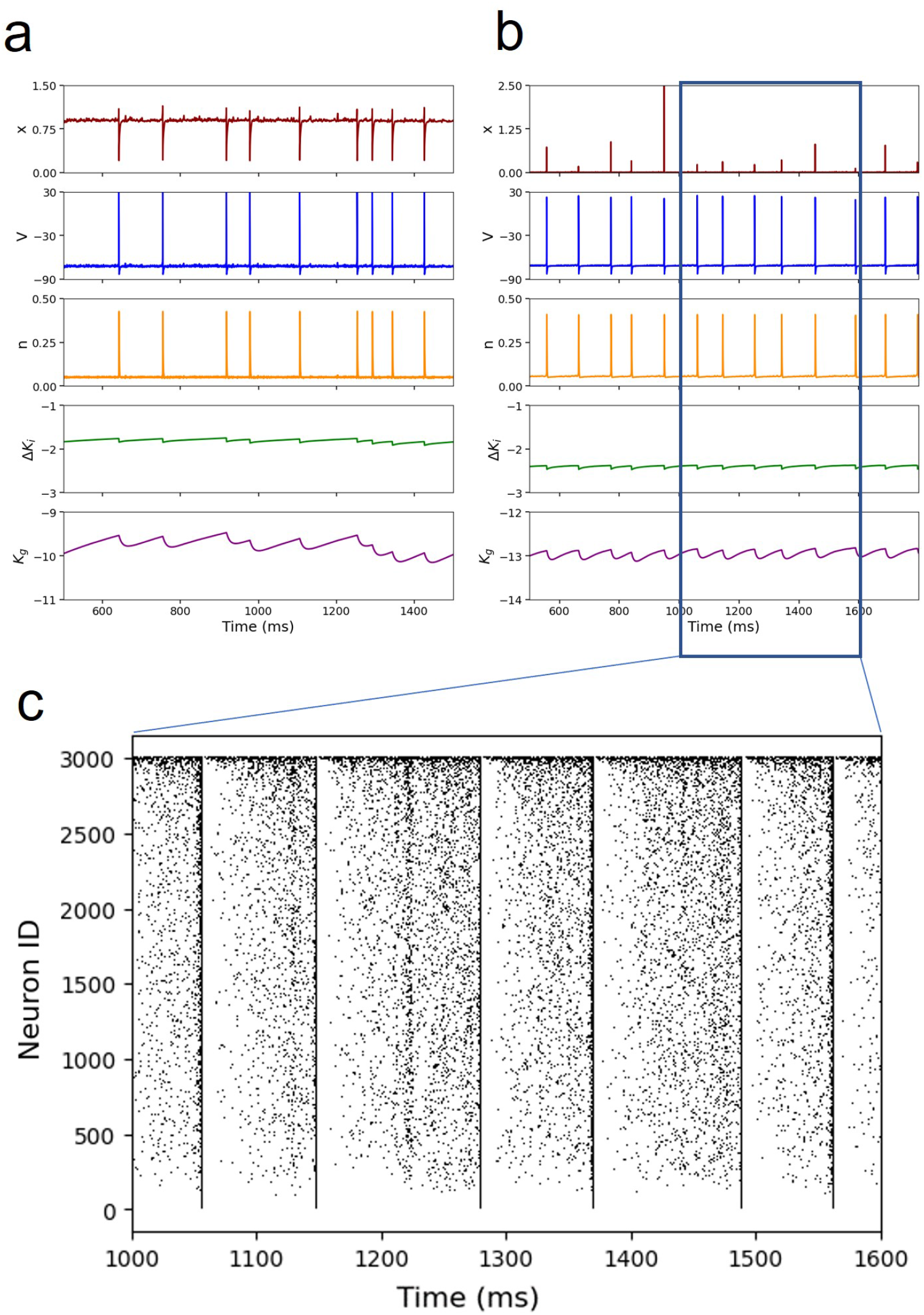
Noise-induced synchronized activity in the asynchronous regime. (a) Time series of *x, V, n*, Δ*K*_*i*_ and *K*_*g*_ for the mean-field model with additive stochastic input, showing intermittent synchronized population events emerging from an otherwise asynchronous state. (b) Corresponding time series for the spiking neural network under matched noise conditions, reproducing the same qualitative alternation between asynchronous and synchronized activity. (c) Raster plot of the spiking network over the time window 1000-1600 ms highlighted in (b), showing that the synchronized events arise from an otherwise irregular, asynchronous firing pattern.

These events are therefore consistent with noise-driven excursions from the excitable resting state rather than transitions between two deterministic equilibria. These results motivate testing the excitable regime as an operating point for future network simulations. Whether it reproduces empirical resting-state activity will depend additionally on structural connectivity, inter-regional coupling, delays and the statistics of the fluctuations, none of which are addressed by the present single-node analysis.

## 3 Discussion

The central distinction in this study is between conditional attractors of a frozen fast subsystem and equilibria of the complete slow-fast model. Holding the ionic variables fixed reveals a broad region with coexisting low- and high-activity fast states, but a full-system equilibrium must also satisfy the slow ionic balance equations. On the physiological resting branch, imposing those conditions leaves a single equilibrium over the range examined, and that state is excitable. This distinction matters for resting-state modeling because a bistable portrait obtained after freezing slow physiology cannot automatically be interpreted as two states available to the complete system. Slow variables are not only parameters that move the fast subsystem through a bifurcation diagram; once dynamical, they also participate in the stability conditions that define the attractors.

Cases 5 and 6 illustrate two different ways in which the frozen map can fail to predict the complete dynamics. In Case 5, the slow variables do not settle but circulate on a limit cycle, so a stationary fast-subsystem label cannot describe the full orbit. In Case 6, the terminal slow-variable values lie in the fast-subsystem bistable region, yet the low fast fixed point fails the slow equilibrium condition and is therefore not a second equilibrium of the complete system. The generic calculation clarifies the same distinction at the level of local stability: for the extracellular sign of the linearized coupling, simultaneous stability of the outer branches requires a coupling above the threshold derived in the *SI Appendix*, whereas for the intracellular sign the two stability intervals do not overlap for positive *λ*_0_ and *λ*_1_. After *K*_*g*_ equilibrates in the KIonEx-MF model, *K*_*o*_ is fixed by *K*_bath_ and Δ*K*_*i*_ remains the relevant ionic degree of freedom in the resting-branch equilibrium calculation.

The spiking-network comparison also shows where the mean-field description is reliable and where it is less accurate. Four regimes—asynchronous activity, synchronized firing, epileptic-like activity, and depolarization block—show close correspondence between the two descriptions. Depolarization-recovery and rapid synchronized firing show larger quantitative differences in voltage range, timing, or population-peak statistics. In both of these cases the slow variables evolve substantially during the activity, making a description based on a single frozen fast subsystem less informative.

In the physiological low-activity state, sufficiently large perturbations produce pronounced excursions before the trajectory returns to the same fixed point. With stochastic input, these excursions appear as intermittent population events separated by longer periods near rest. The spiking network shows the same qualitative pattern, with irregular background firing interrupted by brief periods of stronger population coordination. This provides a mechanism for spontaneous events without deterministic switching between two stable equilibria and motivates testing the excitable state as a node-level operating point in structurally connected brain networks.

Ion-based population models are particularly useful for studying this type of interaction because the slow physiological variables are part of the dynamics rather than external control parameters. The KIonEx-MF formulation belongs to this class, as does Epileptor-2, which incorporates extracellular potassium and intracellular sodium dynamics [15]. Although such models have been used extensively to study pathological activity, the issue considered here is broader. The same difficulty arises whenever the local state space of a neural model is identified while a slow physiological variable that later feeds back onto the dynamics is held fixed.

There are several limitations to the present analysis. We have studied an isolated node, so the results do not rule out bistability generated through coupling between brain regions, delays, or heterogeneous inputs. The depolarization-recovery and rapid synchronized regimes also show larger differences between the mean-field and spiking models than the other four regimes and would benefit from a more detailed analysis of the evolving slow variables. The KIonEx-specific equilibrium elimination is exact for fixed-point existence, but the numerical root count presented here focuses on the physiological resting branch and is not a full continuation of the depolarized branch; moreover, stability and local bifurcations require the full five-dimensional Jacobian.

A natural next step is to embed the excitable regime in a structurally connected whole-brain model and compare the resulting activity with empirical resting-state statistics. The same biophysical node could also be used in pathological network models, including patient-specific epilepsy frameworks such as the Virtual Epileptic Patient [7**? ?**]. The regimes characterized here may also provide summary statistics for parameter-estimation and simulation-based inference approaches [43–45]. More generally, slow physiological variables that feed back onto fast neural dynamics should be included in the stability analysis rather than treated only as fixed parameters of a precomputed bifurcation diagram.

## 4 Materials and Methods

### Mean-field model

The model is a five-dimensional mean-field reduction coupling the Montbrió-Pazo-Roxin description of a population of Hodgkin-Huxley-type neurons [33] to slow potassium exchange dynamics [42]. The state is (*x, V, n*, Δ*K*_*i*_, *K*_*g*_), where *x* is the transformed firing-rate variable of the mean-field reduction, *V* is the mean membrane potential of the population, *n* is a mean gating variable, Δ*K*_*i*_ is the deviation of intracellular potassium concentration from baseline, and *K*_*g*_ tracks extracellular potassium relative to the bath. The equations are

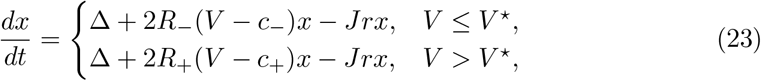

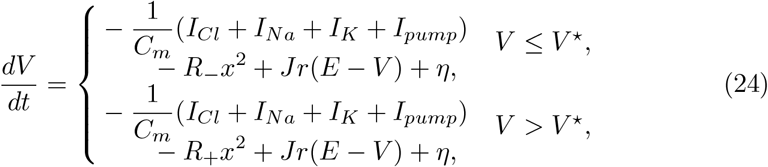

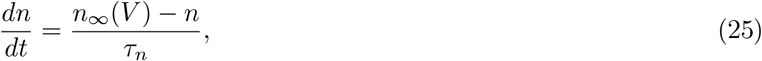

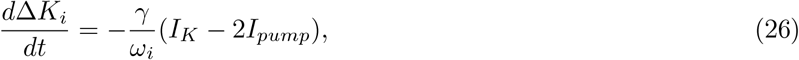

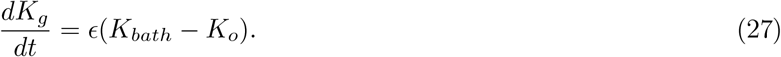

with population firing rate *r* = *R*_*−*_*x/π*, and the ionic currents given by

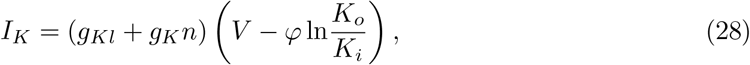

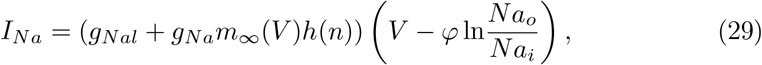

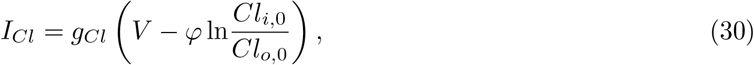

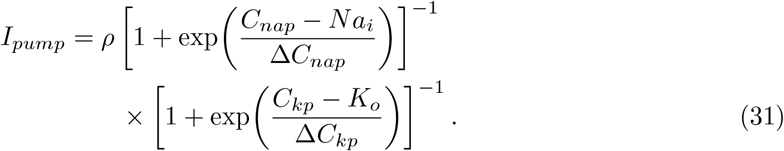

where *φ* = *RT/F* = 26.64 mV, 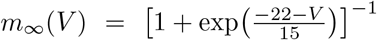, 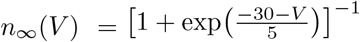, and *h*(*n*) = 1.1 [1 + exp(8(*n* 0.4))]^*−*1^. The intracellular and extracellular ion concentrations entering these currents are themselves determined by the slow state,

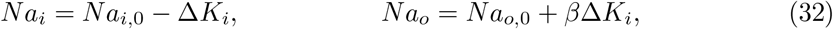

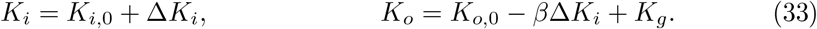

with *β* = *ω*_*i*_*/ω*_*o*_ the ratio of intracellular to extracellular volume. The “Failure” class in Fig. 4 corresponds to parameter combinations for which *K*_*o*_ or *Na*_*o*_ is driven nonpositive by this relationship, making the logarithmic terms undefined.

### Parameter Values

Table 1 lists the parameter values used throughout, unless mentioned otherwise.

**Table 1.**
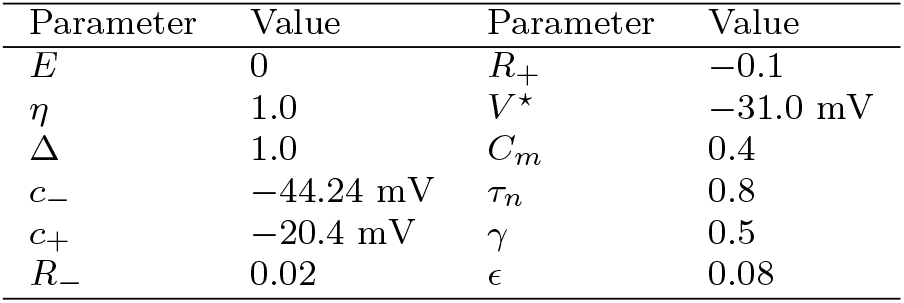
Fixed parameters of the mean-field model.

### Numerical integration

All simulations were performed by numerical integration of the model equations using `scipy.integrate.solve ivp`, with relative and absolute tolerances of 10^*−*8^ and 10^*−*10^, respectively.

### Fast-subsystem classification

For the sweep in Fig. 4, Δ*K*_*i*_ and *K*_*g*_ were fixed at each grid point and the fast subsystem (*x, V, n*) was integrated forward from three initial conditions per point, chosen to sample distinct basins of attraction. Each resulting trajectory was summarized by the mean and peak-to-peak amplitude of *V* over its second half (to exclude the initial transient); a trajectory was classified as a fixed point if this amplitude fell below 0.2 mV, and further split into low- or high-activity by whether the mean *V* fell below or above − 60 mV. Trajectories not classified as fixed points were split into small- and large-amplitude limit cycles by a 20 mV amplitude threshold. The three per-point trajectories were then combined into one of the nine classes of Fig. 4 according to whether they agreed (single fixed point or single limit-cycle class) or disagreed (coexistence of two or more classes, i.e. multistability) with one another, with disagreement in mean *V* or amplitude of at least 0.5 mV/0.5 mV required before two trajectories were considered dynamically distinct rather than numerically noisy repetitions of the same attractor.

### Noise implementation

Stochastic fluctuations were introduced into the *V* and *n* equations as additive Gaussian noise, *dV* → *dV* + *σ*_*A*_ *dW*_*V*_, *dn* → *dn* + *σ*_*A*_ *dW*_*n*_, and separately as multiplicative Gaussian noise scaled by the instantaneous state, *dV* → *dV* + *σ*_*M*_ |*V* − *E*| *dW*_*V*_, *dn* →*dn* + *σ*_*M*_ |*n*| *dW*_*n*_, where *dW*_*V*_ and *dW*_*n*_ are independent Wiener increments.The stochastic equations were integrated using the Euler-Maruyama method. The recovery variables Δ*K*_*i*_ and *K*_*g*_ were left deterministic. The noise-amplitude sweeps in *SI Appendix*, Fig. S6 span *σ*_*A*_ ∈ [0, 0.5] for additive noise and *σ*_*M*_ ∈ [0, 0.05] for multiplicative noise.

### Spiking neural network

Simulations of the spiking network were performed in Brian2 [46]. The network comprised *N* = 3,000 neurons, each governed by the same ionic currents (*I*_*Na*_, *I*_*K*_, *I*_*Cl*_, *I*_*pump*_) and gating-variable dynamics as the mean-field reduction, integrated with RK4 at a fixed time step of 0.0005 ms for 60 s. Neurons were coupled all-to-all, excluding self-connections, through the instantaneous rule 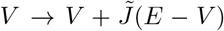 after each presynaptic spike, with 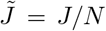. This scaling makes the summed recurrent input comparable to the mean-field term *Jr*(*E* −*V*). A spike was registered when *V* crossed upward through −50 mV. No explicit voltage reset was imposed; a refractory condition prevented repeated detection while *V* remained above threshold, while the membrane potential continued to evolve under the ionic currents.

Heterogeneity across neurons was introduced as in the exact mean-field reduction [33]: a per-neuron constant *η*_*i*_ was drawn from a Cauchy (Lorentzian) distribution with scale Δ centered on 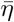, entering the voltage equation as *η*_*i*_*/τ*_*I*_ with *τ*_*I*_ = 1 ms. Draws falling outside 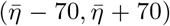 (roughly 1% of the sample, given the heavy Cauchy tail) were replaced with 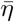 itself rather than redrawn, a small departure from an exact Lorentzian that the mean-field reduction formally assumes. Parameters are the same as in the MF model with the expected *J* → *J/N* modification.

### Data, Materials, and Software Availability

Simulation code and analysis scripts are available upon acceptance.

## Author contributions

S.C., C.M., S.P., H.W., D.D. and V.J. designed research; S.C. and C.M. performed research and analyzed data; D.D. and V.J. supervised this research; S.C. wrote the original draft; C.M., S.P., D.D., H.W. and V.J. reviewed and edited the manuscript.*D.D. and V.J. are co-last authors

---

**Table S1.** Comparison of voltage statistics between the mean-field (MF) model and the spiking neural network (SNN). All reported *V* values are in mV. The final column reports detected peaks/events in the population-averaged voltage signal; it does not count individual neuronal spikes.

| Regime | Model | $V_{\min}$ | $V_{\text{mean}}$ | $V_{\max}$ | $V_{\text{sd}}$ | Population peaks |
| --- | --- | --- | --- | --- | --- | --- |
| Asynchronous | MF | -70.28 | -70.28 | -70.28 | 0.00 | 0 |
|  | SNN | -70.65 | -70.38 | -69.86 | 0.11 | 0 |
| Synchronized firing | MF | -71.91 | -57.41 | 27.03 | 27.41 | 223 |
|  | SNN | -72.69 | -57.38 | 29.97 | 28.58 | 249 |
| Epileptic-like activity | MF | -78.22 | -52.45 | 26.94 | 29.83 | 32 |
|  | SNN | -78.07 | -52.08 | 24.71 | 30.17 | 35 |
| Depolarization-recovery | MF | -87.36 | -51.07 | 25.97 | 30.20 | 0 |
|  | SNN | -45.46 | -24.47 | -1.29 | 7.11 | 0 |
| Rapid synchronized firing | MF | -30.38 | -23.57 | -16.06 | 4.97 | 1954 |
|  | SNN | -56.67 | -29.62 | 10.23 | 22.18 | 785 |
| Depolarization block | MF | -21.76 | -21.76 | -21.76 | 0.00 | 0 |
|  | SNN | -22.31 | -22.26 | -22.20 | 0.02 | 0 |

## Supplementary Materials

### Parameter-Sweep Statistics

Figure S1 shows the six summary statistics of the membrane-potential time series from which the regime boundaries of Fig. 1 (main text)a were derived. Each panel is itself a (*K*_*bath*_, *J*) sweep, computed over the same 1000 ms window used throughout:standard deviation and (b) median of *V* separate quiescent, low-variance regions from oscillatory ones and locate the depolarized-block plateau; (c) minimum and (d) maximum of *V* delineate the voltage range spanned in each regime, (e) the number of detected population-voltage peaks and (f) the mean population-voltage peak height jointly separate regimes that share similar voltage statistics but differ in whether population events are present. No single statistic in isolation separates all six regimes; the boundaries in Fig. 1 (main text)a were obtained by combining thresholds on several of these panels simultaneously.

### Quantitative comparison between the mean-field and spiking network dynamics

To quantify agreement and discrepancies between the two descriptions, we compared the population-averaged membrane potential generated by the mean-field (MF) model with the average membrane potential of the spiking neural network (SNN) for the six representative dynamical regimes shown in Fig. 1(b). For each regime, summary statistics were computed over a 1-s window (or over one representative activity cycle for periodic epileptic-like and depolarization-recovery states).

Table S1 summarizes the principal voltage statistics for both models. For the asynchronous, synchronized firing, epileptic-like, and depolarization block regimes (Cases 1, 2, 3 and 6), the mean-field model closely reproduces the collective dynamics of the spiking network. In the asynchronous regime, both models remain near −70 mV with negligible fluctuations and no detected events in the population-averaged voltage signal (individual neurons in the SNN continue to fire). During synchronized population firing, the two descriptions also have similar voltage extrema and mean voltages and produce comparable population-peak counts (223 in the MF model and 249 in the SNN).

**Fig. S1.**
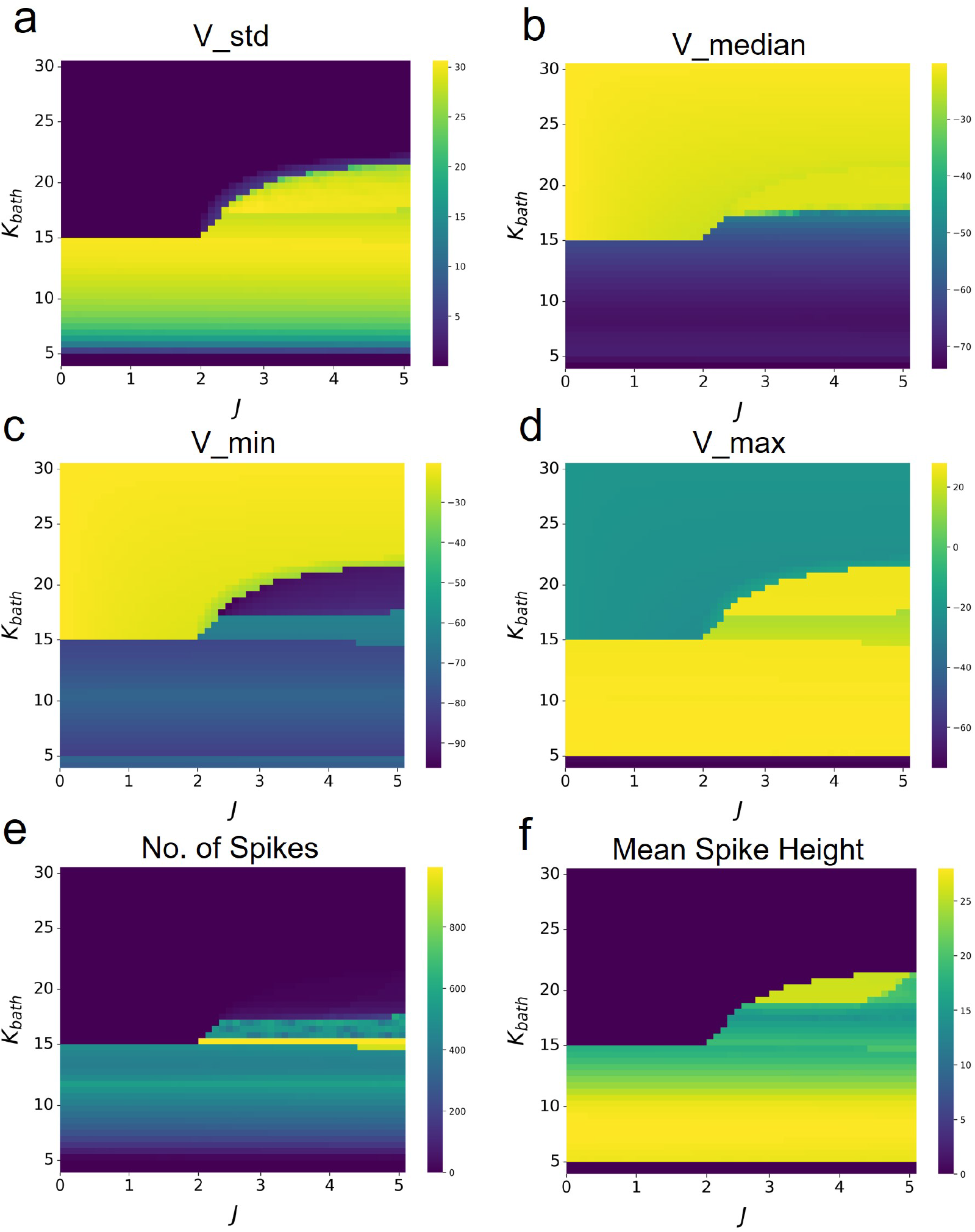
Membrane-potential statistics used to define the regime boundaries in Fig. 1 (main text)a. For each point in the (*K*_*bath*_, *J*) parameter sweep, statistics of *V* were computed over a 1000 ms window (1500-2500 ms) of simulation: (a) standard deviation, (b) median, (c) minimum, (d) maximum, (e) number of detected population-voltage peaks, and (f) mean peak height. Thresholds applied to these statistics were combined to define the six regime boundaries shown in red in Fig. 1 (main text)a.

Agreement is similarly observed during the epileptic-like regime. Over one representative episode, the voltage extrema, mean membrane potential, and temporal variability are similar in the two models. The mean-field model produces 32 population-voltage peaks during an episode lasting approximately 91.9 ms, whereas the spiking network produces 35 peaks over approximately 96.2 ms. Thus, for this representative case, the reduced model captures the principal timescale and voltage range of the population event.

In the depolarization-block regime, both models converge to a stable depolarized equilibrium near −22 mV with negligible temporal variance and no detected population-voltage events. This regime therefore shows close quantitative correspondence between the mean-field and spiking descriptions.

Larger quantitative differences appear in the depolarization-recovery and rapid synchronized firing regimes (Cases 4 and 5). During depolarization-recovery, the meanfield model predicts a larger excursion between hyperpolarized and depolarized phases, whereas the population-averaged voltage of the spiking network exhibits substantially smaller excursions due to asynchronous recruitment and incomplete synchronization of individual neurons. Similarly, in the rapid synchronized regime, the event detector identifies 1954 population-voltage peaks in the 1-s mean-field analysis window and 785 peaks in the corresponding spiking-network window. The SNN oscillations are broader because of microscopic variability and partial synchrony.

To examine these two regimes in more detail, Fig. S2 shows the spiking-network raster plot underlying the depolarization-recovery regime (Case 4), and Fig. S3 compares the slow-variable limit cycle traced out by Δ*K*_*i*_ and *K*_*g*_ in the mean-field model against the spiking network for the rapid synchronized regime (Case 5).

The raster in Fig. S2 shows population recruitment during the depolarized epochs and reduced activity during recovery. The two descriptions differ in both voltage excursion and timing: the SNN has a substantially narrower population-averaged voltage range (Table S1), and its depolarized and recovery epochs are less regular in duration than the near-periodic mean-field alternation.

The larger slow-variable loop in the spiking network is consistent with the broader population-voltage excursions reported for Case 5 in Table S1.

### Epileptic-like and Depolarization-recovery activity

In the main text we noted that the epileptic-like (Case 3) and depolarization-recovery (Case 4) regimes cannot be described by a single stationary point of the fast subsystem, because the membrane potential alternates between two qualitatively distinct activity states over the course of a cycle. Figure S4 makes this alternation explicit. For both cases, we split one representative cycle into its two constituent phases and color-code them light red and light green in the *V*, Δ*K*_*i*_ and *K*_*g*_ time series (panels a, d).

Overlaying the same two-colored trajectory on the fast-subsystem nullclines (panels b, e) shows that each phase of the cycle occupies a distinct region of the fast phase space, separated by a transition where the trajectory departs the neighborhood of one set of nullcline intersections and is captured by another; the crosses mark these transitions. Panels c and f re-plot the same trajectory in reduced coordinates to trace out the full hysteresis loop directly. The two cases differ in the detailed geometry of this loop - consistent with the different shape of their *V* time series in Fig. 1 (main text)b - but share the same underlying picture: it is the slow drift of Δ*K*_*i*_ and *K*_*g*_ that carries the fast subsystem from one branch to the other, rather than either branch being a stable terminus of the dynamics on its own.

### Slow-Variable Trajectories Overlaid on the Classification Map

Figure S5 overlays the trajectory traced by Δ*K*_*i*_(*t*) and *K*_*g*_(*t*) on the frozen fast-subsystem classification map of Fig. 4 (main text) for each representative case. The star and triangle mark the initial and terminal slow-variable values. Cases 5 and 6 show why the frozen classification is conditional. In Case 5 (panel e), the slow variables continue to circulate while the instantaneous fast subsystem lies in the low-fixed-point region, so the full system remains oscillatory. In Case 6 (panel f), the terminal slow-variable values lie in the frozen bistable region, but evaluating the slow balance on the low fast-subsystem branch gives Δ*K*_*i*_ ≠0; that low branch is therefore not a fixed point of the complete system. Across the initial conditions tested, the full system converged to the high-activity state.

### Noise-Amplitude Dependence of Synchronized-Event Frequency

Figure S6 summarizes how the number of noise-triggered synchronized events, counted in the asynchronous regime of Fig. 5 (main text), scales with noise amplitude for (a) additive and (b) multiplicative noise. Both noise types show the same qualitative shape: synchronized events are rare or entirely absent below a threshold noise amplitude, and beyond that threshold the event count grows approximately linearly. The similarity between the additive and multiplicative cases suggests that this threshold behavior is a property of the excitable geometry identified in Fig. 3 (main text).

### Loss of bistability in the presence of slow ionic dynamics

#### Cusp reduction of the exact QIF mean field

We begin from the exact Montbrió–Pazó–Roxin mean-field equations for a heterogeneous population of quadratic integrate-and-fire neurons [33]. The firing rate *r* and mean membrane potential *v* obey

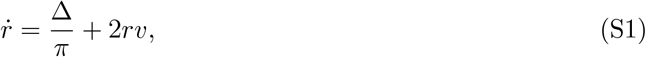

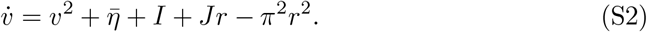

Introducing *w* = *πr* + *iv* combines these two equations into

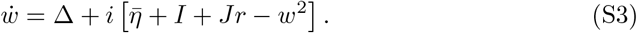

At equilibrium, 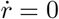 gives *r* = −Δ*/*(2*πv*). Substitution into 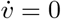 yields the quartic

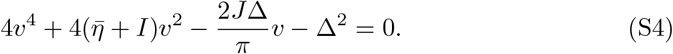

The two saddle-node curves bounding the bistable region meet where this polynomial has a triple root. Setting the polynomial and its first two derivatives to zero gives, for *I* = 0,

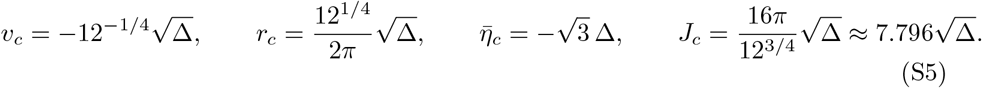

The Jacobian at the cusp has eigenvalues {0, 4*v*_*c*_}, so the local dynamics has one center direction and one contracting direction. A center-manifold reduction therefore gives,to cubic order,

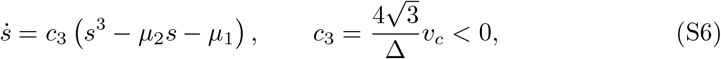

which is the unfolded cusp used in the main text after rescaling time by |*c*_3_ |. The outer roots are attracting and the middle root is repelling within the bistable wedge.

#### Generic fast–slow stability calculation

For the calculation, we shift the slow coordinate by its reference value, *y* = *u* − *u*_0_, and rescale the cusp normal form so that

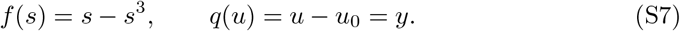

The linearized potassium-current contribution is written

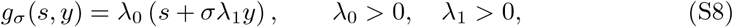

where *λ*_0_ = *g*_*s*_ is the dimensionless coupling of the current to the fast coordinate after rescaling, and *λ*_1_ fixes the magnitude of its slow-coordinate derivative. The sign *σ* = −1 corresponds to the extracellular case (*g*_*u*_ *<* 0), whereas *σ* = +1 corresponds to the intracellular case (*g*_*u*_ *>* 0). The coupled system is

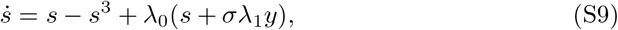

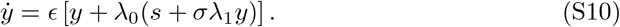

At a fixed point, the slow equation gives

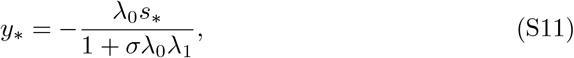

and the fast equation, together with *g*_*σ*_ = −*y*_*\**_ at equilibrium, gives 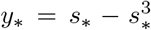. Hence the nonzero outer equilibria satisfy

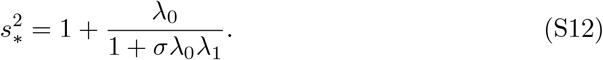

The Jacobian is

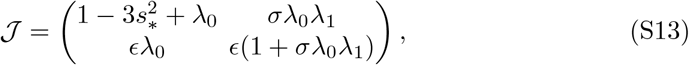

with

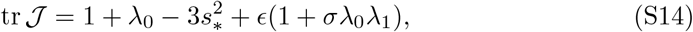

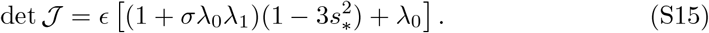

After eliminating *u* to obtain a scalar fixed-point equation *H*(*s*) = 0, the same determinant can be written as det *J* = *ϵ*(*q*^*′*^ + *g*_*u*_)*H*^*′*^(*s*_*\**_).

For extracellular coupling (*σ* = −1), define *c* = *λ*_0_*λ*_1_. Equation (S12) becomes

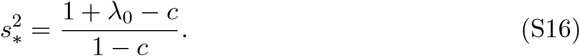

At this outer equilibrium,

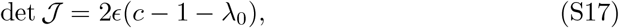

so positive determinant requires *c >* 1 + *λ*_0_. In the singular limit *ϵ* →0, the leading trace is

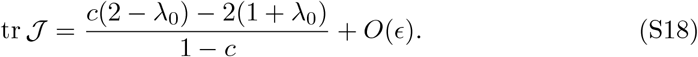

Because 1− *c <* 0 once the determinant condition is satisfied, the trace is negative when

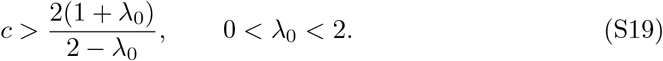

This condition is stronger than *c >* 1 + *λ*_0_ and therefore gives

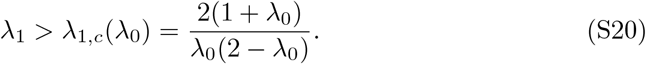

Differentiating the right-hand side with respect to *λ*_0_ gives its minimum at 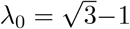,where

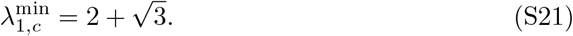

For finite but small *ϵ*, the exact trace condition differs from Eq. (S20) by an *O*(*ϵ*) correction; the threshold quoted in the main text is the slow-fast limit.

For intracellular coupling (*σ* = +1), stability requires, to leading order in *ϵ*,

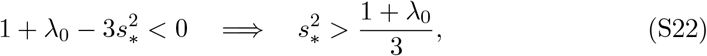

whereas det *J >* 0 requires

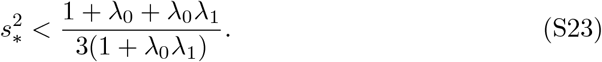

The two intervals could overlap only if

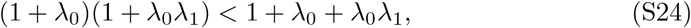

which reduces to

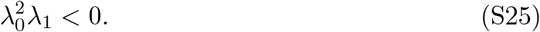

This is impossible for *λ*_0_, *λ*_1_ *>* 0, so the two outer branches cannot both be stable in this linearized intracellular case.

#### Local oscillation timescale near the fold

Two small quantities enter the local dynamics near a fold. The ionic equation contributes the explicit slow factor *ϵ*, while the eigenvalue of the cusp-reduced fast dynamics, *λ*_c_ = *c*_3_(3*s*^2^ −*µ*_2_), approaches zero at the fold. Away from the fold, the eigenvalues of the coupled fast–slow system separate into an order-one branch and an order-*ϵ* branch. In the local scaling used for the generic model, the leading-order balance at the fold gives

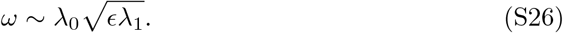

Thus the small-oscillation frequency is set by the geometric mean of the relevant fast and slow rates. The square-root dependence is compatible with singular-Hopf asymptotics; identifying the local bifurcation itself additionally requires the corresponding eigenvalue and nondegeneracy conditions.

#### Exact equilibrium elimination of the firing rate and gating variable

For equilibrium counting, setting 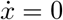 and 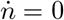 is exact rather than adiabatic. Because the model defines *r* = *R*_*−*_*x/π* on both voltage branches, 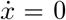 gives the branchwise quadratic

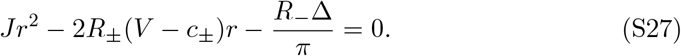

On the resting branch, the physical positive root is

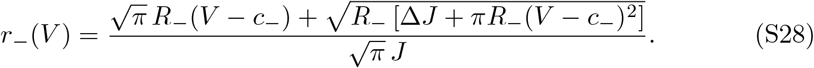

On the depolarized branch,

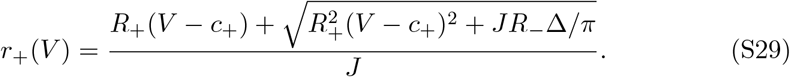

The quantity under the square root is strictly positive for *J*, Δ, *R*_*−*_ *>* 0. Because the square root exceeds |*R*_+_(*V* −*c*_+_)|, Eq. (S29) gives one positive and one negative quadratic root for every real *V* ; the physical firing-rate branch is therefore real-valued throughout the depolarized branch. At equilibrium, *n* = *n*_*∞*_(*V*) exactly.

#### Closed-form currents after elimination

After imposing *n* = *n*_*∞*_(*V*) and the *K*_*g*_ equilibrium condition *K*_*o*_ = *K*_*bath*_, the currents needed for the reduced (*V*, Δ*K*_*i*_) system can be written as

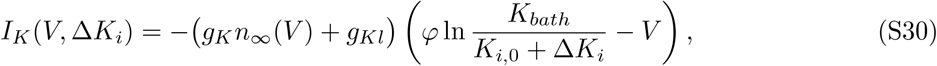

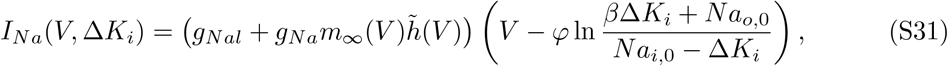

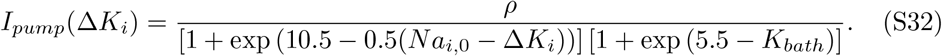

Here 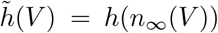. The chloride current has no Δ*K*_*i*_ dependence under the assumptions of the model, and after *K*_*o*_ = *K*_*bath*_ the pump depends on Δ*K*_*i*_ through the intracellular sodium term but has no explicit *V* dependence.

#### Root counting and scope of the equilibrium reduction

At a full-system equilibrium, 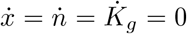 implies Eqs. (S27)–(S29), *n* = *n*_*∞*_(*V*), and *K*_*o*_ = *K*_*bath*_ exactly. Substituting these relations into 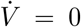 and Δ*K*_*i*_ = 0 therefore gives an exact two-variable algebraic problem for equilibrium existence and location. On the physiological resting branch, the Δ*K*_*i*_-nullcline *F* (*V*, Δ*K*_*i*_) = 0 was solved for Δ*K*_*i*_ over the physical bracket (−*K*_*i*,0_, *Na*_*i*,0_) ≈ (−130, 16) using `scipy.optimize.brentq`.The resulting curve Δ*K*_*i*_ = *f* (*V*) was substituted into the voltage nullcline 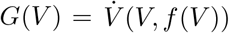, and its zero crossings were counted for each *K*_*bath*_ tested. Exactly one crossing was found over the resting branch examined in the main text.

Equation (S29) shows that the firing-rate relation remains real on the depolarized branch; there is no narrow discriminant-limited voltage interval. The present numerical root count was performed on the physiological resting branch, so we do not use it as an exhaustive continuation of depolarized equilibria. When both branch equations are enumerated, however, the algebraic reduction itself gives the exact fixed-point count. Linear stability is a separate question and must be evaluated from the full five-dimensional Jacobian.

#### Nonlinear check of the intracellular-feedback case

The linearized intracellular-potassium case in the main text has no parameter range with two simultaneously stable outer branches for *λ*_0_, *λ*_1_ *>* 0. Direct integration of the corresponding linearized two-variable model produces unbounded growth rather than convergence, but that behavior is partly a consequence of replacing the Nernst logarithm by an unsaturated linear coupling. Replacing the linearized current with the full logarithmic Nernst form keeps the fast variable bounded in this minimal test. The slow variable can still grow because the simplified intrinsic term *q*(*u*) = *u*− *u*_0_ contains no additional saturating restoration. We therefore use this calculation only as a check of the feedback sign and stability condition, not as a global model of ionic concentration dynamics.

**Fig. S2.**
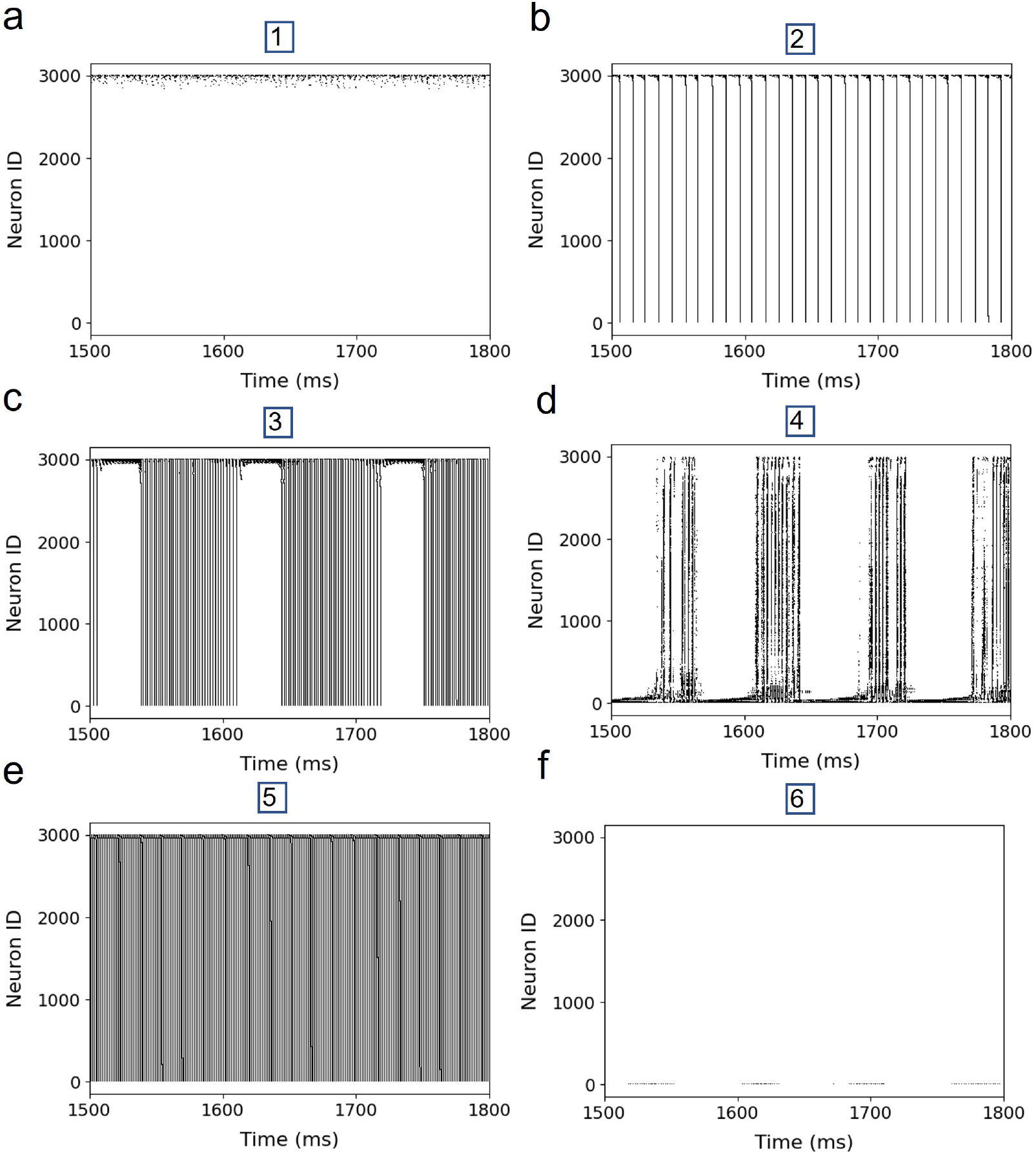
Spiking-network raster plots for the six representative dynamical regimes. (a) Case 1, asynchronous activity. (b) Case 2, synchronized population firing. (c) Case 3, epileptic-like activity. (d) Case 4, depolarization-recovery activity. (e) Case 5, rapid synchronized firing. (f) Case 6, depolarization block. Each panel shows a 300 ms representative window (1500-1800 ms) of spiking activity across all 3,000 neurons in the network, qualitatively matching the corresponding mean-field dynamics of Fig. 1 (main text)b.

**Fig. S3.**
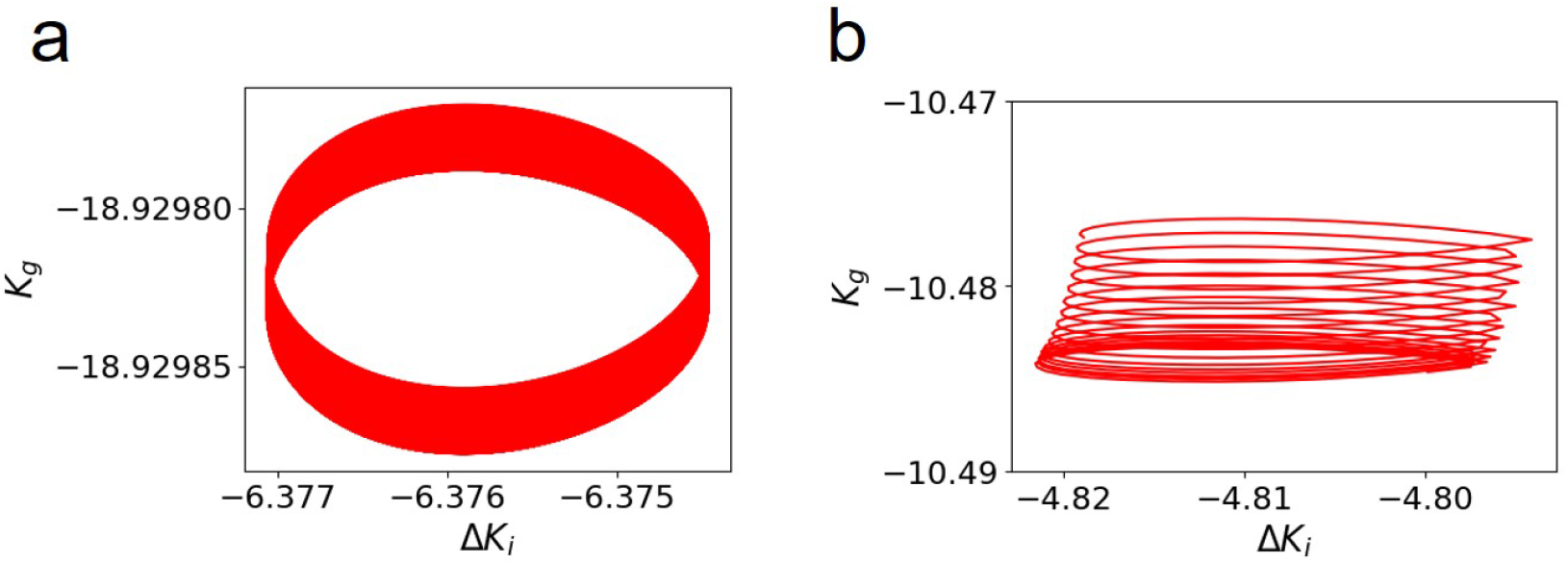
Slow-variable limit cycles for Case 5 (rapid synchronized firing). Trajectories in the (Δ*K*_*i*_, *K*_*g*_) plane for (a) the mean-field model and (b) the spiking neural network. The SNN limit cycle spans a substantially larger range of both Δ*K*_*i*_ and *K*_*g*_ than the mean-field model, mirroring the larger oscillation amplitude of *V* observed for this regime in the spiking network (cf. Table S1).

**Fig. S4.**
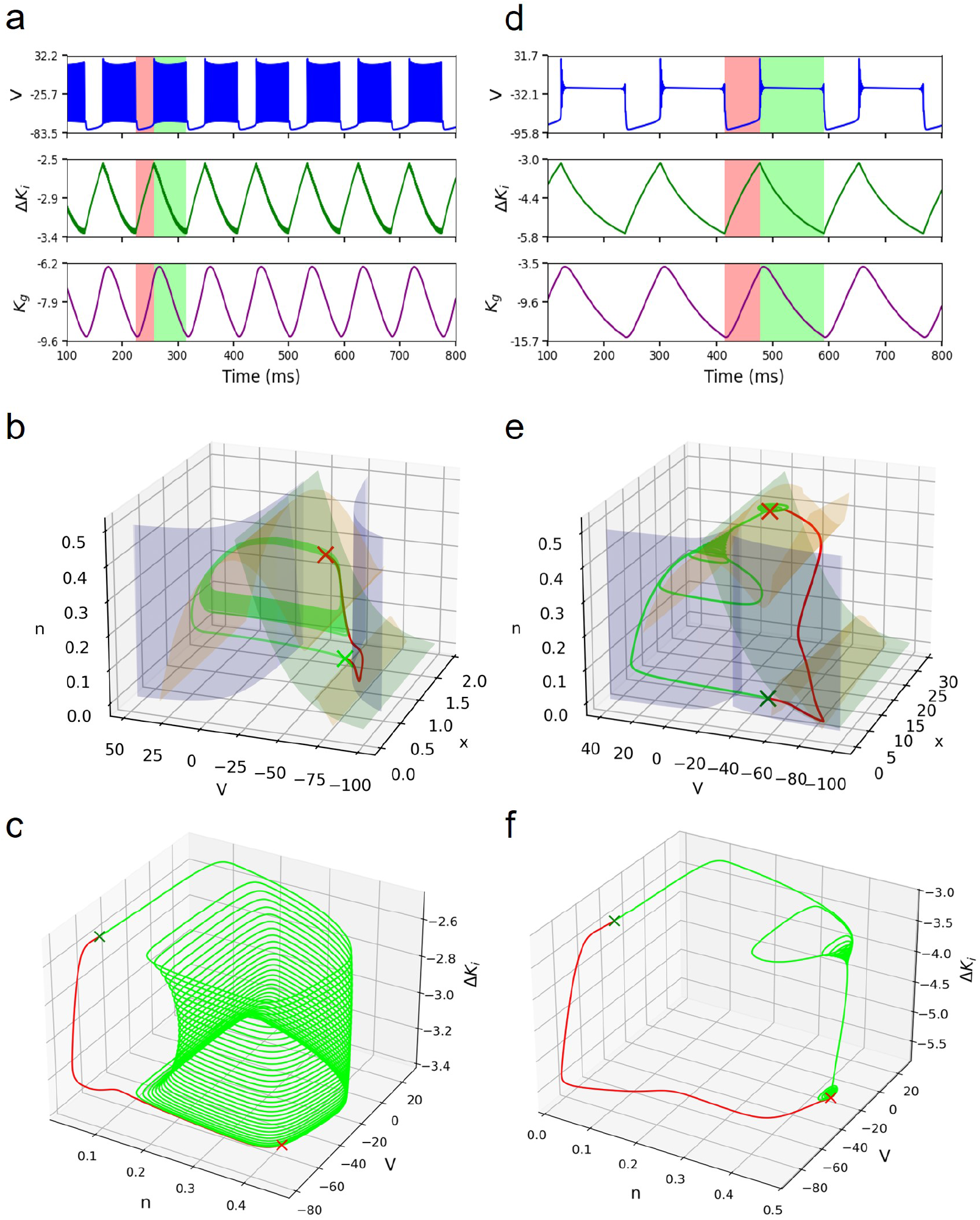
Hysteresis underlying the two-state dynamics of Case 3 (epileptic-like activity, left, a-c) and Case 4 (depolarization-recovery activity, right, d-f). (a, d) Time series of *V*, Δ*K*_*i*_ and *K*_*g*_, with the two constituent activity states of each cycle color-coded in light red and light green. (b, e) The corresponding fast-subsystem trajectory, colored to match (a, d), overlaid on the fast-subsystem nullclines (as in Fig. 2 (main text)); crosses mark the transitions between the two states. (c, f) The same trajectory shown in (*V, n*, Δ*K*_*i*_) space, illustrating the hysteresis loop traced as the fast subsystem alternates between the two activity states.

**Fig. S5.**
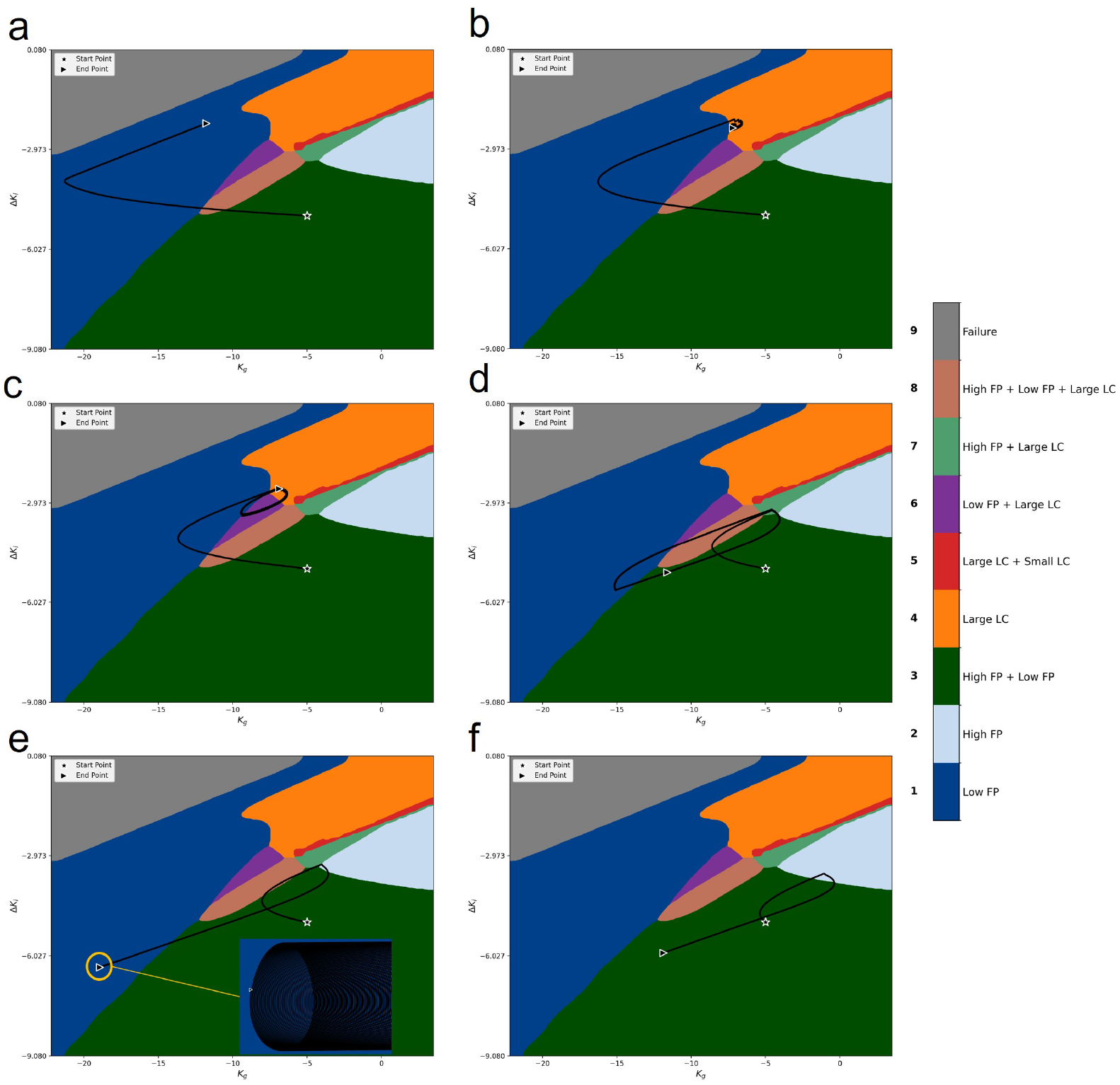
Slow-variable trajectories of the full system for the six representative cases, overlaid on the fast-subsystem classification map of Fig. 4 (main text). Panels (a-f) correspond to Cases 1-6 of Fig. 1 (main text), respectively; the star and triangle mark the start and end point of each trajectory in (Δ*K*_*i*_, *K*_*g*_) space. Comparing the frozen classification with the full-system dynamics (Fig. 1 (main text)b) shows where the two descriptions differ. In (e), the slow variables circulate while the frozen subsystem lies in a low-fixed-point region, and the complete system remains oscillatory. In (f), the final slow-variable values lie in the frozen bistable region, but the low fast-subsystem branch does not satisfy the slow equilibrium condition and therefore is not a second fixed point of the complete system. **Abbreviations:** FP, fixed point (Low, low-activity; High, high-activity); LP, limit cycle (Small, small-amplitude; Large, large-amplitude).

**Fig. S6.**
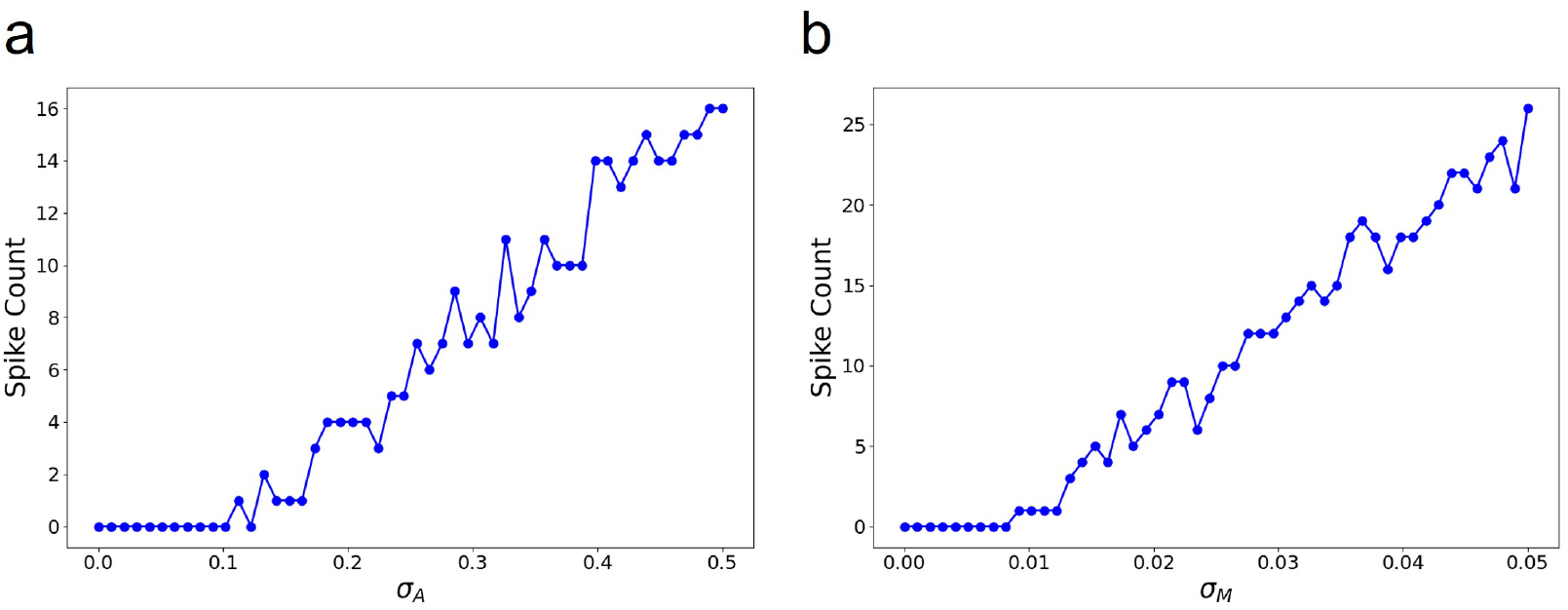
Dependence of noise-induced synchronized-event frequency on noise amplitude, in the asynchronous regime. Number of synchronized population events as a function of (a) additive noise strength *σ*_*A*_ and (b) multiplicative noise strength *σ*_*M*_. In both cases, synchronized events are rare or absent below a threshold noise level, beyond which event frequency increases approximately linearly with noise amplitude.

## Notes

### Competing Interest Statement

The authors have declared no competing interest.

### Funding

- Horizon europe: 101147319, 101057429

